# Characterization of Genetic and Phenotypic Heterogeneity of Obstructive Sleep Apnea Using Electronic Health Records

**DOI:** 10.1101/724443

**Authors:** Olivia J. Veatch, Christopher R. Bauer, Navya Josyula, Diego R. Mazzotti, Brendan T. Keenan, Kanika Bagai, Beth A. Malow, Janet D. Robishaw, Allan I. Pack, Sarah A. Pendergrass

**Author notes:** Corresponding Authors: Olivia J. Veatch, MS PhD, Research Associate, 125 S. 31^st^ St, Office 2123, Philadelphia, PA 19104,; Sarah A. Pendergrass, PhD MS, Assistant Professor, Biomedical and Translational Informatics Institute, Geisinger Research, 6101 Executive Blv, Suite 110, North Bethesda, MD 20852. **Author Contributions:** AIP, JDR, & SAP conceived of the presented idea. OJV, CRB, DRM, BTK & SAP designed the analysis plan with critical feedback from JDR, KB, BAM & AIP. OJV, DRM & BTK conducted the literature review. OJV, CRB & NJ conducted all analyses. OJV drafted the initial manuscript. All authors discussed contributed to the final manuscript.

## Abstract

Obstructive sleep apnea (OSA) is defined by frequent episodes of reduced or complete cessation of airflow during sleep and is linked to negative health outcomes. Understanding the genetic factors influencing expression of OSA may lead to new treatment strategies. Electronic health records can be leveraged to both validate previously reported OSA-associated genomic variation and detect novel relationships between these variants and comorbidities. We identified candidate single nucleotide polymorphisms (SNPs) via systematic literature review of existing research. Using datasets available at Geisinger (n=39,407) and Vanderbilt University Medical Center (n=24,084), we evaluated associations between 48 SNPs and OSA diagnosis, defined using clinical codes. We also evaluated associations between these SNPs and OSA severity measures obtained from sleep reports at Geisinger (n=6,571). Finally, we used a phenome-wide approach to perform discovery and replication analyses testing associations between OSA candidate SNPs and other clinical codes and laboratory values. Ten SNPs were associated with OSA diagnosis in at least one dataset, and one additional SNP was associated following meta-analysis across all datasets. Three other SNPs were solely associated in subgroups defined by established risk factors (i.e., age, sex, and BMI). Five OSA diagnosis-associated SNPs, and 16 additional SNPs, were associated with OSA severity measures. SNPs associated with OSA diagnosis were also associated with codes reflecting cardiovascular disease, diabetes, celiac disease, peripheral nerve disorders and genitourinary symptoms. Results highlight robust OSA-associated SNPs, and provide evidence of convergent mechanisms influencing risk for co-occurring conditions. This knowledge can lead to more personalized treatments for OSA and related comorbidities.

## INTRODUCTION

In the United States (US), 50-70 million individuals suffer from a disorder of sleep and wakefulness^1^. Among the most common sleep disorders is obstructive sleep apnea (OSA). OSA is defined by frequent episodes of reduced (hypopnea) or complete cessation (apnea) of airflow that occur as a consequence of upper airway obstruction during sleep. Approximately 34% of men and 17% of women aged 30-70 exhibit mild OSA, and 13% and 6%, respectively, suffer from moderate to severe OSA^2^. Notably, prevalence rates substantially increase by the age of 50 years old^2^. When left untreated, OSA represents a significant public health burden carrying a higher risk of serious comorbidities such as cardiovascular disease^3–11^, cancer incidence^12^ and mortality, all causes^13; 14^, cognitive impairment^15^, and rate of progression of neurodegeneration^16^. As such, identifying more effective diagnosis and management of OSA are important areas of research.

OSA is a complex, phenotypically heterogeneous disorder driven by both obesity-related factors and other mechanisms^17–19^. A number of studies have observed that OSA has a heritable component, with a two-fold increased risk in first-degree relatives of individuals with OSA^20–26^. Genomic variation is also strongly implicated in some of the well-established structural risk factors for OSA (e.g., soft tissue volumes^27^, craniofacial dimensions^28–31^, and obesity^32–34^). Given the high heritability of obesity and the strong relationship with OSA, it is important to note that studies indicate OSA familial aggregation is not simply due to association with obesity^21; 24^. Genomic variation is also associated with measures of OSA severity, like the apnea-hypopnea index (AHI)^19^. Furthermore, a number of syndromes with well-defined genetic causes are associated with increased prevalence of OSA (e.g., Achondroplasia^35^, Down syndrome^36; 37^, Marfan’s syndrome^38^, Prader-Willi syndrome^39^). Understanding the underlying molecular mechanisms that contribute to expression of OSA, and to distinct patient subgroups, may offer the opportunity to inform novel, more personalized approaches to treatment.

Despite the substantial evidence implicating genetic factors on expression of OSA, no strong genetic candidates have been robustly established^19; 40^. This is likely due to underpowered genetic association studies^19; 40^, lack of independent replication^41; 42^, and the substantial heterogeneity in the expression of OSA limiting the ability to identify genomic variants affecting all patients similarly. With the expansion of biorepositories linked to electronic health records (EHR), large sample sizes and a variety of OSA-related information—as well as other clinical and diagnostic data—are available to evaluate candidate genes and variants previously implicated in OSA. These resources offer unprecedented opportunity to establish strong genetic risk factors contributing to OSA, understand factors contributing to risk of OSA in various population subgroups, and detect potential pleiotropic genetic effects contributing to comorbidities in OSA. Several genes currently implicated in OSA are associated with co-occurring conditions, suggesting shared etiologies^2; 34; 43–49^. Phenome-wide association studies (PheWAS), which test candidate variants for associations with a broad range of phenotypes^50–52^, is a powerful approach to help reveal pleiotropic genetic effects.

In this study, we use EHR data linked to genomic data to better understand OSA etiology and heterogeneity, as well as the overlapping genetic mechanisms influencing expression of co-occurring conditions. We first determined if previously reported associations between a comprehensive list of candidate single nucleotide polymorphisms (SNPs) and OSA validated in clinical samples. We then used PheWAS to provide evidence of pleiotropic effects for OSA-associated SNPs. This work represents one of the largest candidate gene studies, and first PheWAS, focused on OSA.

## METHODS

### Search strategy and selection criteria for OSA-associated genes and variants

PubMed and PubMed Central (PMC) were queried for original research papers and previous reviews published up to 08/31/2017. Search terms were as follows: (obstructive sleep apnea[Title/Abstract] OR obstructive sleep apnoea[Title/Abstract])) AND (polymorphism OR genetic variant OR genetic association OR gene OR genome wide association study OR genome-wide association study). PubMed results were filtered to only include studies conducted in humans (i.e. adults and children reflecting all ancestral backgrounds) and manuscripts written in English. Manuscripts were excluded if there was evidence of being retracted. Abstracts were manually reviewed to ensure that the results related to genetic studies conducted in humans that were specifically relevant to OSA as opposed to phenotypes that may reflect something other than an OSA diagnosis (e.g., intermittent hypoxia, sleep-disordered breathing, treatment response, obesity). Studies were further excluded if there were no full-texts available, or represented analyses of larger genomic regions for which individual variants could not be identified (e.g., linkage analysis). The remaining eligible full manuscripts were manually reviewed to identify candidate variants with evidence for association with OSA. Candidate variants were then mapped to rsIDs and independent SNPs (r^2^<0.50) were selected using genome-wide data (described in more detail below) from individuals in the Geisinger dataset. Data were linkage disequilibrium pruned, while adjusting for cryptic relatedness. When possible, proxy SNPs (r^2^≥0.50) for candidate OSA variants were identified using data from 1000 Genomes, phase III^53^ Europeans using the rAggr web tool (http://raggr.usc.edu/).

### Genotype quality control and imputation

SNPs that were previously reported to be associated with OSA and OSA severity were tested for association with EHR-derived OSA and other EHR-derived traits in three independent datasets, including individuals whose data were available in clinic-based biorepositories linked to EHRs from two sites (i.e., Geisinger European Americans, Vanderbilt University Medical Center (VUMC) European Americans, and VUMC African Americans). Genotype data for the Geisinger dataset were generated using the Illumina® HumanOmniExpressExome bead chip. Genotype data for the VUMC datasets were generated using the Illumina® Multi-Ethnic Global Array (MEGA). Quality control (QC) procedures required DNA samples to have >90% genotyping call rate and be unrelated based on PI-HAT≤0.05. Directly genotyped markers were required to have >99% call rate with minor allele frequency >0.01, and p-values testing significant deviation from Hardy-Weinberg Equilibrium >1×10^−7^. Genetically informed ancestry was then determined using principal components analysis with reference human genomes available via the 1000 Genomes Project, phase III^53^. The QCd genotype data for the European ancestral dataset from Geisinger were imputed using Impute2 software and reference human haplotypes available via the Haplotype Reference Consortium^54^. The QCd genotype data for the European and African VUMC ancestral datasets were imputed using Impute2 and reference human haplotypes available via the 1000 Genomes Project, phase III^53^. Imputed genotypes were required to have info scores ≥0.3. Notably, while all other QC thresholds were required for inclusion in association tests, given the goal to specifically evaluate previously reported OSA candidate SNPs for associations with EHR-derived OSA status, some imputed SNPs with >1% missingness in the VUMC datasets were tested for associations with an OSA diagnosis. Specifically, there were six imputed SNPs that were missing in >1% of the VUMC European ancestral dataset: rs1801278 (missing=22.7%), rs1799983 (missing=15.2%), rs1054135 (missing=41.8%), rs9526240 (missing=24.6%), rs9932581 (missing=79.7%), rs2743173 (missing=15.9%). In addition, there were five imputed SNPs that were missing in >1% of the VUMC African ancestral dataset: (rs999944 (missing=19.3%), rs1801278 (missing=14.1%), rs7804372 (missing=10.5%), rs9526240 (missing=10.9%), rs2743173 (missing=16.0%)).

### Validation of previously reported OSA-associated genomic variants using diagnoses in electronic health records

EHR-derived OSA cases (EHR-OSA) and non-cases (NonOSA) were defined using an algorithm based on the total instances of International Classification of Diseases 9^th^ (ICD-9) or 10^th^ (ICD-10) revision OSA-related diagnostic codes (Table S1). This algorithm was validated with clinical chart reviews at Geisinger and VUMC^55^. Based on the EHR algorithm validation study, EHR-derived OSA was defined at each site using the minimum number of code instances that achieved positive predictive value (PPV) and negative predictive value (NPV) of at least 90%. For the Geisinger dataset, EHR-OSA was defined as individuals with any OSA-related codes present on at least three different dates in the EHR (PPV [95% CI] =94.9 [88.5, 98.3]; NPV [95% CI] = 95.0 [88.7, 98.4])^55^. At VUMC, EHR-OSA was defined as individuals with any OSA-related codes present on at least two different dates in the EHR (PPV = 97.5 [92.9, 99.5]; NPV = 94.0 [87.4, 97.8])^55^. Non-cases were defined as individuals with zero OSA-related diagnostic codes in the EHR. Individuals not meeting the case or non-case definitions (i.e., with exactly one code instance at VUMC or exactly one or two instances at Geisinger) were excluded. The final Geisinger analysis dataset was comprised of 39,407 individuals (n_OSA_=5,760, n_NonOSA_=33,647) with genetically informed European ancestry. The Geisinger sample is approximately 97% European American and due to the sample sizes of other ancestral groups being limited^56^, these were not included in this study. The final VUMC analysis sets included 20,688 individuals (n_OSA_=2,831, n_NonOSA_=17,857) with European ancestry and 3,396 individuals (n_OSA_=331, n_NonOSA_=3,065) with African ancestry. Sample sizes for populations from VUMC with other genetically informed ancestries were too small to evaluate (total n=328 across all populations; data not shown), and therefore were excluded from this study.

The associations between SNPs previously implicated in OSA, that were selected during literature reviews, and the validated EHR-derived diagnosis of OSA were evaluated using logistic regression models, adjusting for age, sex and body mass index (BMI). For individuals with EHR-OSA, age and BMI measures reflected those at the time of the first usage of an OSA code, for individuals with no evidence of OSA the most recent age and BMI measures were included in analyses. SNPs were coded additively with respect to the minor allele (0, 1 or 2 copies). As the purpose of this study was to validate individual candidate SNPs, each with *a priori* evidence of effect, our significance was set at an uncorrected p<5.0×10^−2^. Results from EHR-OSA association tests across the three datasets were then meta-analyzed using fixed-effects inverse-variance weighted meta-analyses in the METAL software package^57^. The possibility of heterogeneity among the studies was also tested using METAL.

To determine if age, sex or BMI modified the associations between any candidate SNP and EHR-OSA, we performed statistical interaction tests by including a product term (SNP x Covariate) in the logistic models along with main effect terms. If this product term was significant (p<5.0×10^−2^), we tested the associations between OSA candidate SNPs and EHR-OSA separately in subgroups stratified by the covariate of interest, adjusted for age, BMI and sex (except when performing analyses in males/females separately). Age groups were defined as ≤50 years and >50 years, based on evidence that prevalence of OSA increases substantially above the age of 50 years-old^2^. BMI groups were stratified as ≤30 kg/m^2^ and BMI>30 kg/m^2^, based on accepted US standards of distinguishing normal-overweight from obese^58^.

### Tests for associations between previously reported OSA candidate variants and quantitative traits obtained from sleep study reports at Geisinger

Sleep reports linked with genome-wide data were available for a subset of the analysis dataset from Geisinger (n_NonOSA_=1,614, n_EHR-OSA_=4,957); these data were not available for the VUMC datasets. Linear regression models were used to test for additive effects of the minor allele at OSA candidate SNPs and 12 quantitative traits pulled from sleep study (i.e. polysomnography [PSG]) reports included in the EHR, adjusting for age, sex and BMI (defined based on EHR-OSA status as described above). The focus was on respiratory indices (Percent Time SaO2<89% [during Non-Rapid Eye Movement or Rapid Eye Movement], Respiratory Event Duration, Minimum Percent of SpO2 [during Respiratory Event or Total Sleep Time], Apnea/Hypopnea Index, Central Apnea Index, Duration of Apnea/Hypopnea) and measures of sleep quality (Number of Awakenings, Sleep Efficiency, Wake After Sleep Onset), as well as patient-responses to the Epworth Sleepiness Scale (ESS)^59^. Only results obtained from diagnostic PSG with ≥120 minutes of total sleep time were included in analyses. If an individual had >1 sleep report, variables reflect results from the diagnostic sleep study that was conducted on the date closest to the first usage of the OSA code. For individuals with no evidence of OSA in the EHR, variables were pulled from the most recent sleep study performed. For the majority of the quantitative sleep report traits, the distributions did not meet the assumptions of normality necessary for regression analyses. Prior to analyses, Box-Cox power transformations were applied to sleep study variables and then Z-scores were calculated. For variables that had zero values, half of the minimum non-zero value was added to each observation.

### Phenome-wide association tests for previously reported OSA candidate variants and other EHR-derived clinical traits: discovery (Geisinger) and replication (VUMC)

Logistic regression models were used to test for associations between the previously reported OSA-associated SNPs and other non-OSA clinical codes, while adjusting for age, sex and BMI (defined based on EHR-OSA status as described above). For discovery analyses, only those codes resulting in a minimum sample size of 200 cases in the Geisinger dataset were included. For each code being tested, ‘cases’ were defined as individuals with codes used on ≥3 different dates in the EHR and ‘controls’ were defined as individuals with absence of this code or a code within the same hierarchy. Individuals with one or two instances of a code were excluded from the association analysis for that code. Linear regression models were used to test for associations between OSA candidate SNPs and clinical laboratory values. For discovery analyses, only laboratory values available for ≥1,000 individuals in the Geisinger dataset were included. In total, OSA candidate SNPs were tested for associations with 577 non-OSA ICD codes and 143 different quantitative traits defined using Logical Observation Identifier Names and Codes (LOINC) mapped median clinical laboratory values. Box-Cox power transformations were applied for normality following inspection of the laboratory value distributions.

Codes and laboratory values that were associated with any candidate SNP based on a false discovery rate multiple testing correction threshold (q<5.0×10^−2^) were tested for replication in the VUMC European ancestral dataset and generalization in the VUMC African ancestral dataset. The significance threshold for replication and generalization was set at an uncorrected p<5.0×10^−2^. To help circumvent issues related to potential differences in specific ICD code usage across the two clinical sites, all ICD codes that reflected the same phenotype code (PheCode) as the discovery ICD code, present on ≥3 different dates in the individual’s record, were tested for replication in the VUMC datasets. PheCodes are aggregations of one or more related ICD code into distinct diseases or traits^60^. Prior to PheCode mapping, all ICD-10 codes were mapped to the corresponding ICD-9 code using CMS General Equivalency Mapping available via the Agency for Healthcare Research and Quality Map It Tool, Application Version 5.1.110, data version 2.2018.110X. ICD-9 codes and converted ICD-10 codes were then mapped to the corresponding PheCode based on PheCode Map 1.2^60^. In addition, for OSA candidate SNPs that were associated with EHR-OSA, full PheWAS were run in the corresponding dataset. For SNPs associated with EHR-derived OSA in specific covariate-defined subgroups, full PheWAS were run separately in the corresponding strata.

## RESULTS

### Literature-derived identification of OSA candidate variants

A systematic literature review of 204 studies revealed 53 unique SNPs reported to be associated with OSA and/or OSA severity (Figure S1). Of the 53 SNPs, we were able to evaluate associations with 48 variants: 46 were directly assessed, proxy SNPs were identified for two (literature-derived SNP: rs35424364, proxy SNP: rs7752028, r^2^=0.97; literature-derived SNP: rs25531, proxy SNP: rs11080123, r^2^=0.50), and five SNPs were dropped due to low allele frequencies (MAF<0.01). Details for the 48 selected candidate SNPs, including the reported effects and hyperlinks to the literature sources, are provided in Table S2. As discussed above, analyses were performed to validate these literature-reported associations in three independent datasets representing clinical samples from two different US-based sites: Geisinger (European American) and VUMC (European American and African American).

### Sample Characteristics

Among the Geisinger sample of European ancestry, the mean age was 59 years (range 19-88 years), 41% of participants were male, the mean BMI was 31.61 kg/m^2^ (range: 14.17-89.06, kg/m^2^), and 14.6% of individuals met the EHR criteria for OSA. Individuals with EHR-defined OSA were older, had higher BMIs and were more likely to be male than non-OSA participants (Table 1). Among the VUMC European ancestry sample, the mean age was 57 years (range: 18-88 years), 42.4% were male and the mean BMI was 28.80 kg/m^2^ (range: 10.71-93.65 kg/m^2^). A total of 13.7% of individuals met the criteria for EHR-defined OSA, and OSA participants were younger, had higher BMIs and were more likely to be male (Table 1). Among the VUMC African ancestral sample, the mean age was 47 years (range: 18-88 years), 37.8% were male and the mean BMI was 31.00 kg/m^2^ (range: 10.59-78.73 kg/m^2^). OSA was defined using EHR-criteria in 9.7% of individuals. These individuals had higher BMIs compared to EHR-defined NonOSA; however, age and sex were not significantly different at p<5.0×10^−2^ (Table 1).

**Table 1.** Demographic Characteristics of the Analysis Datasets.

| Analysis Sample | Demographics | EHR-defined No OSA | EHR-defined OSA | p-value<sup>#</sup> |
| --- | --- | --- | --- | --- |
| Geisinger<br>(European Ancestry) | N | 33,647 | 5,760 | — |
| | Age<sup>£</sup>, years | 58.9 (17.2) | 61.1 (12.6) | $2.65 \times 10^{-29}$ |
| | BMI<sup>£</sup>, kg/m<sup>2</sup> | 30.1 (6.8) | 40.1 (9.8) | $2.20 \times 10^{-16}$ |
| | Male, % | 38.9% | 52.9% | $2.15 \times 10^{-87}$ |
| Vanderbilt University<br>Medical Center<br>(European Ancestry) | N | 17,857 | 2,831 | — |
| | Age<sup>£</sup>, years | 56.8 (18.2) | 55.8 (13.0) | $2.88 \times 10^{-4}$ |
| | BMI<sup>£</sup>, kg/m<sup>2</sup> | 27.9 (6.6) | 34.3 (9.0) | $1.08 \times 10^{-241}$ |
| | Male, % | 40.1% | 56.9% | $1.93 \times 10^{-62}$ |
| Vanderbilt University<br>Medical Center<br>(African Ancestry) | N | 3,065 | 331 | — |
| | Age<sup>£</sup>, years | 47.3 (17.6) | 49.2 (13.8) | $5.73 \times 10^{-2}$ |
| | BMI<sup>£</sup>, kg/m<sup>2</sup> | 30.2 (8.0) | 38.5 (10.4) | $2.91 \times 10^{-36}$ |
| | Male, % | 37.4% | 41.7% | $1.37 \times 10^{-1}$ |

### Associations of OSA candidate variants with EHR-derived OSA

Eleven of the 48 OSA candidate SNPs tested were associated with an EHR-derived diagnosis of OSA in at least on dataset (Tables 2 & S2, Figure 1). Specifically, five candidate SNPs were associated with EHR-OSA in individuals of European ancestry in the Geisinger dataset. These included two missense variants (rs1137100, β=−0.08, p=3.58×10^−3^; and rs1137101, β=−0.05, p=3.78×10^−2^) in the *leptin receptor* (*LEPR*) gene, intronic variants in the *lysophosphatidic acid receptor 1* (*LPAR1*) gene (rs7030789, β=0.05, p=3.57×10^−2^) and the *bleomycin hydrolase* (*BLMH*) gene (rs11080123, β=0.13, p=2.18×10^−2^; proxy SNP for the *Solute Carrier Family 6 Member 4* [*SLC6A4*] gene promoter variant (rs25531), and a variant ~2kb upstream of the *matrix metallopeptidase 9* (*MMP9*) gene (rs3918242; β=0.07, p=2.65×10^−2^). Three additional OSA candidate SNPs were associated with EHR-OSA in individuals with European ancestry in the VUMC dataset. These included a missense variant located in the *adrenoceptor beta 2* (*ADRB2*) gene (rs1042713; β=−0.07, p=2.71×10^−2^), a variant ~2kb upstream of the coding region for the *tumor necrosis factor alpha* (*TNF-α*) gene (rs1800629; β=0.10, p=1.31×10^−2^), and a variant ~2kb upstream of the coding region for the *cytochrome b-245 alpha chain* (*CYBA*) gene (rs9932581; β=−0.16, p=3.15×10^−2^). Two additional OSA candidate SNPs were associated with EHR-OSA in individuals of African ancestry in the VUMC dataset. These included an intronic variant in the *neuregulin 1* (*NRG1*) gene (rs10097555; β=0.20, p=2.30×10^−3^), and a variant located in the 3’ untranslated region (3’-UTR) of the *fatty acid binding protein* (*FABP4*) gene (rs1054135; β=−0.30, p=6.03×10^−3^). The meta-analysis which combined results across all three datasets indicated that there was one OSA candidate SNP, a synonymous variant in the *gamma-aminobutyric acid type B receptor subunit 1* (*GABBR1*) gene, that was associated with EHR-OSA among all of the samples (rs29230; β=−0.05, p=2.27×10^−2^, I^2^=0, p_heterogeneity_=9.79×10^−1^).

**Table 2.**
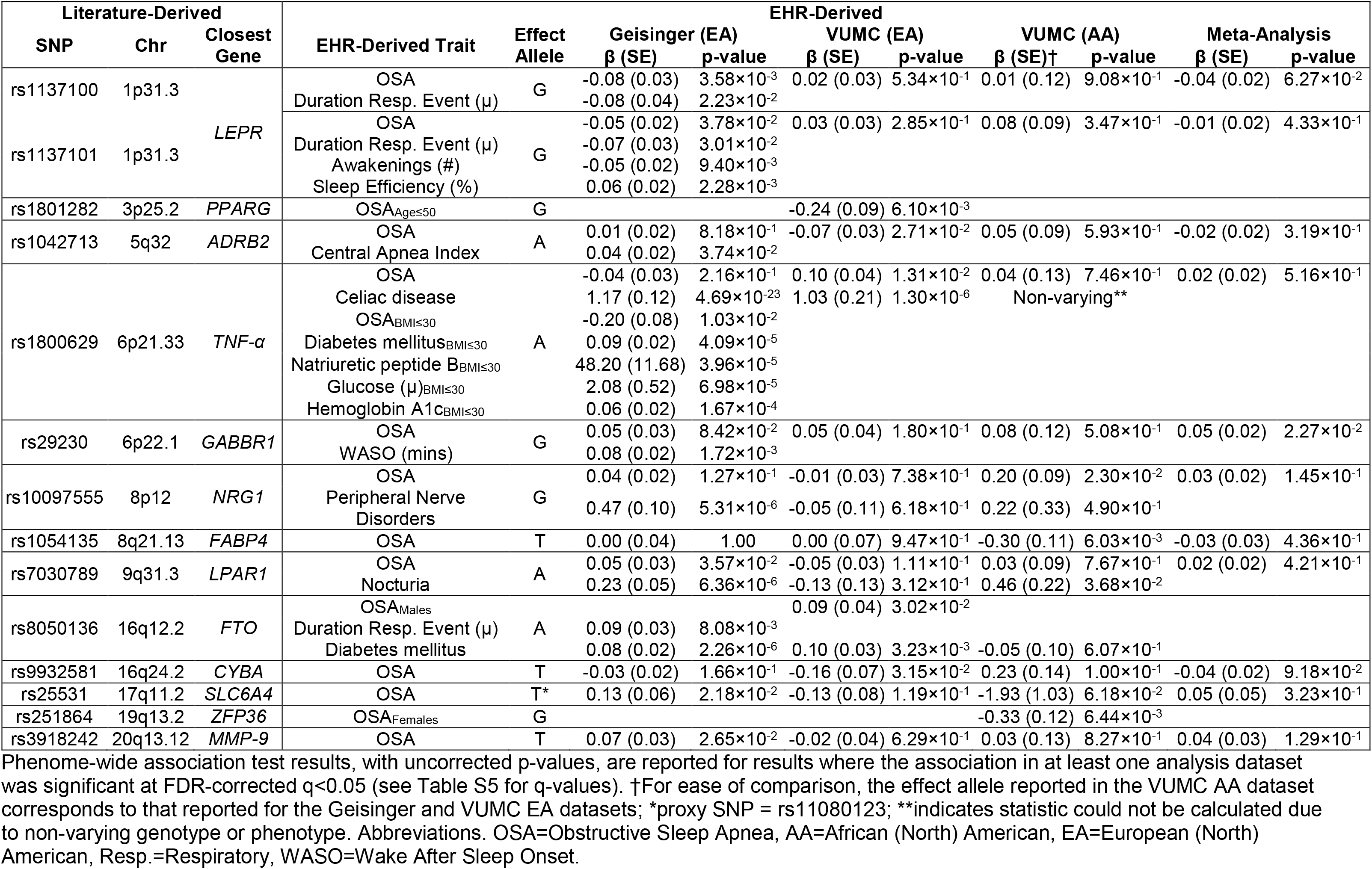
Combined Evidence for SNPs Associated with EHR-Derived Obstructive Sleep Apnea in at Least One Analysis Dataset.

**Figure 1.**
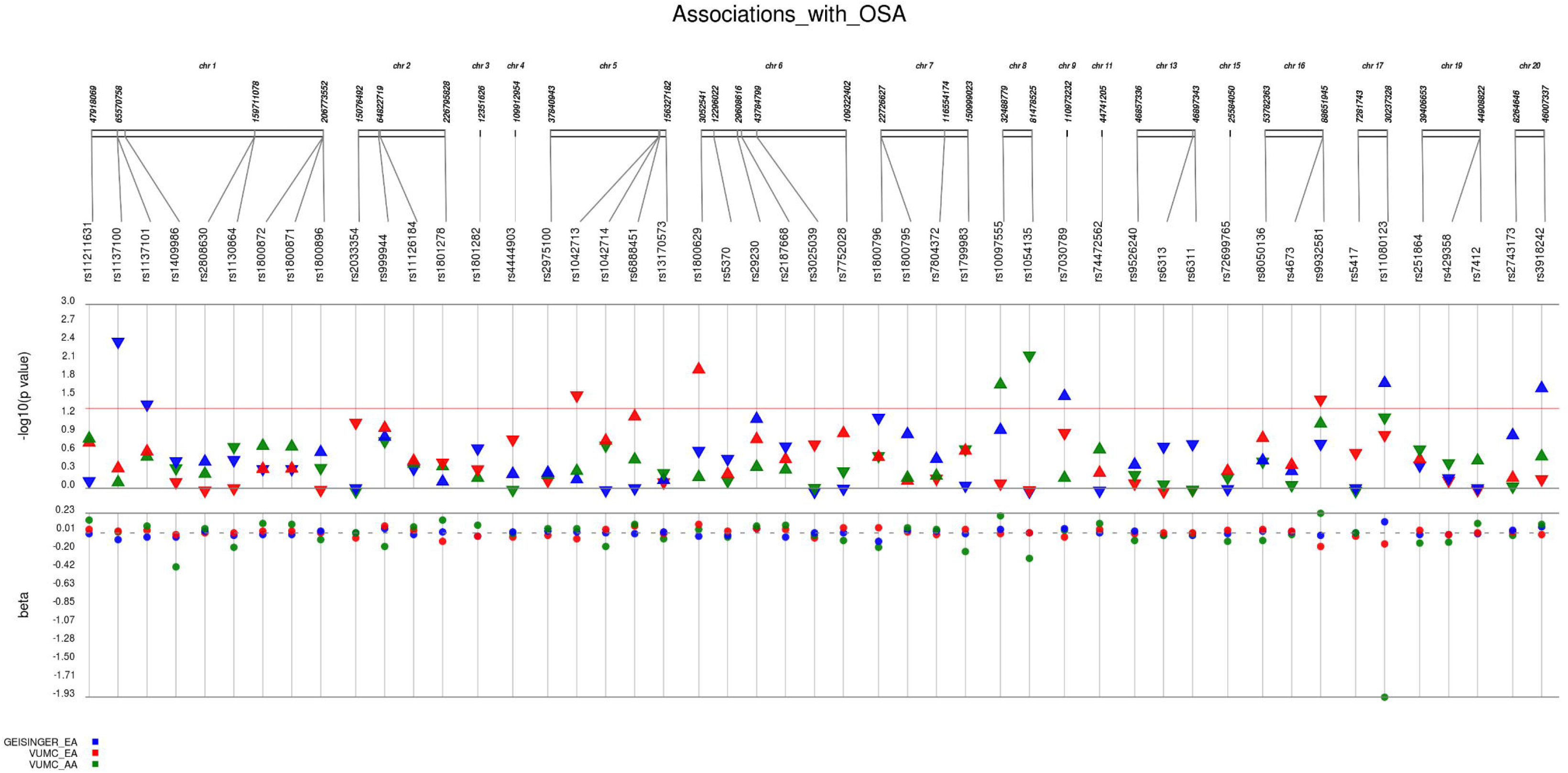
Associations between Literature-derived Candidate SNPs and EHR-derived Obstructive Sleep Apnea. Plotted are the estimates of the additive effects (beta) of the minor allele at each candidate SNP on a validated definition of OSA obtained from the EHR, along with the corresponding −log_10_ p-value. Results from analyses conducted in the Geisinger European American (EA) dataset are plotted in blue, VUMC European Americans in red and VUMC African Americans (AA) in green with the effect adjusted to the EA dataset minor alleles for ease of comparison. For p-values, up arrows denote increased risk for EHR-derived OSA given the minor allele at this SNP and down arrows denote reduced risk.

### Effects of Age, Sex and BMI on Associations of OSA candidate variants with EHR-OSA

The associations between 15 OSA candidate SNPs and EHR-OSA were modified by at least one OSA-related risk factor (i.e., age, sex, and/or BMI) evaluated in interaction tests (Table S3). Interaction tests conducted in the Geisinger dataset indicated that age, sex or BMI modified the associations between seven SNPs and EHR-OSA. However, only one SNP was associated (p<5.0×10^−2^) with EHR-OSA in stratified analyses (Figure S2, Table S3). Specifically, age had an effect on the association between an intronic variant in the *PPARG coactivator 1 beta* (*PPARGC1B*) gene and EHR-OSA (rs6888451; Interaction β=−0.01, p=6.22×10^−3^). No differences were observed in subgroups stratified by clinically relevant values of age (≤50 years [p=6.64×10^−1^] vs. >50 years [p=8.48×10^−1^] years). Similarly, while there was evidence of differential effects in males and females for three SNPs: rs251864 (β=0.21, p=2.68×10^−2^), rs1800872 (β=−0.10, p=3.98×10^−2^) and rs1409986 (β=−0.18, p=9.67×10^−3^), no significant associations were observed with EHR-OSA when examining associations separately in males or females (Table S3). Finally, BMI had an effect on the associations between three SNPs and EHR-OSA: rs1130864 (β=0.01, p=4.48×10^−2^), rs1800629 (β=0.01, p=5.20×10^−3^) and rs7804372 (β=0.01, p=4.11×10^−2^). Analyses conducted separately in normal/overweight (BMI≤30 kg/m^2^) and obese (BMI>30 kg/m^2^) participants indicated that the rs1800629 variant upstream of *TNF-α* was associated with EHR-OSA primarily in individuals with BMI≤30 (β=−0.20, p=1.03×10^−2^) rather than those with BMI>30 (β=−0.01, p=7.69×10^−1^).

In the VUMC European American dataset, age, sex or BMI modified the relationship between eight SNPs and EHR-OSA (Table S3). Of these, two SNPs were associated (p<5.0×10^−2^) with EHR-OSA in covariate-defined subgroups (Figure S2, Table S3). Age modified the association between a missense variant in the *peroxisome proliferator activated receptor gamma* (*PPARG*) gene and EHR-OSA (rs1801282; β=0.01, p=6.64×10^−3^). The association between this SNP and EHR-OSA was primarily observed in individuals who were ≤50 years old (β=−0.24, p=6.10×10^−3^), rather than those who were older than 50 (β=0.06, p=2.54×10^−1^). The associations between two SNPs and EHR-OSA were modified by sex (Table S3). Tests conducted separately in males versus females indicated that the association between an intronic variant in the *fat mass and obesity-associated* (*FTO*) gene (rs8050136; β=−0.14, p=2.91×10^−2^) and EHR-OSA was primarily observed in males (β=0.09, p=3.02×10^−2^), rather than females (β=−0.02, p=6.62×10^−1^). The association between a missense variant in the *apolipoprotein E* (*APOE*) gene and EHR-OSA was also influenced by sex (rs429358; β=−0.22, p=1.66×10^−2^), but was not significantly associated in males or females (Table S3). BMI also influenced the association between the rs429358 SNP in *APOE* with EHR-OSA (β=0.02, p=8.56×10^−3^), as well as modified the effects of five additional SNPs on EHR-OSA (Table S3). However, there were no significant associations when stratifying into normal/overweight and obese participants (Table S3).

In the VUMC African American dataset, age, sex or BMI modified the associations between three SNPs and EHR-OSA (Table S3). Two SNPs were differentially associated with EHR-OSA in covariate-defined subgroups (Figure S2, Table S3). In particular, age modified the association between an intronic variant in the *kinetochore associated 1* (*KNTC1*) gene and EHR-OSA (rs10734924; β=0.02, p=2.16×10^−2^), but no differences were observed when stratifying by age (Table S3). Sex modified the association between a variant located ~2kb upstream of the *ZFP36 ring finger protein* (*ZFP36*) gene and EHR-OSA (rs251864; β=−0.45, p=2.74×10^−2^); this association was observed primarily in females (β=−0.33, p=6.44×10^−3^) but not in males (β=0.19, p=1.89×10^−1^). BMI influenced the association between an intronic variant in the *TraB domain containing 2B* (*TRABD2B*) gene and EHR-OSA (β=−0.03, p=2.20×10^−2^), with the association observed primarily in individuals with BMI>30 (β=−0.25, p=5.40×10^−2^), rather than with BMI≤30 (β=0.15, p=4.75×10^−1^).

### Associations of OSA candidate variants with sleep study report traits

There were 21 OSA candidate SNPs that were associated with at least one quantitative measure of OSA severity obtained from sleep study reports at Geisinger (Figure 2, Tables 2 & S4). Of these, one SNP was the *GABBR1* rs29230 variant which was associated in the meta-analysis across all datasets. Two other SNPs were associated with EHR-OSA in the Geisinger dataset, two in the VUMC European American dataset, and 16 SNPs had no significant evidence for associations with EHR-OSA (Table S4). Notably, the G allele at the *GABBR1* SNP rs29230 was associated with both increased risk for EHR-OSA when results from analyses across all three datasets were meta-analyzed, and increased wake after sleep onset in the subset of the Geisinger dataset with available sleep study reports (β=0.08, p=1.73×10^−3^). In addition, the two missense variants in the *LEPR* gene that were associated with EHR-OSA were also associated with the mean duration of the respiratory event (rs1137100, β=−0.08, p=2.23×10^−2^; rs1137101, β=−0.07, p=3.01×10^−2^, respectively). Futhermore, presence of the A allele at the rs1137101 SNP, which was associated with reduced risk for EHR-OSA, was also associated with a decreased number of awakenings (β=−0.05, p=9.40×10^−3^) and an increased percentage of sleep efficiency (β=0.06, p=2.28×10^−3^). Notably, the intronic *FTO* variant which was associated with increased risk for EHR-OSA solely in males with European ancestry from the VUMC dataset was also associated with a longer mean duration of the respiratory event in the entire Geisinger dataset (rs8050136, β=0.09, p=8.08×10^−3^). There was also a missense variant in *ADRB2* that was associated with reduced risk for EHR-OSA in European Americans from the VUMC dataset but an increased Central Apnea Index in the Geisinger dataset (rs1042713, β=0.04, p=3.74×10^−2^). While not significant, the beta calculated for this SNP in the Geisinger dataset indicated a positive direction of effect for EHR-OSA (β=0.01, p=8.18×10^−1^).

**Figure 2.**
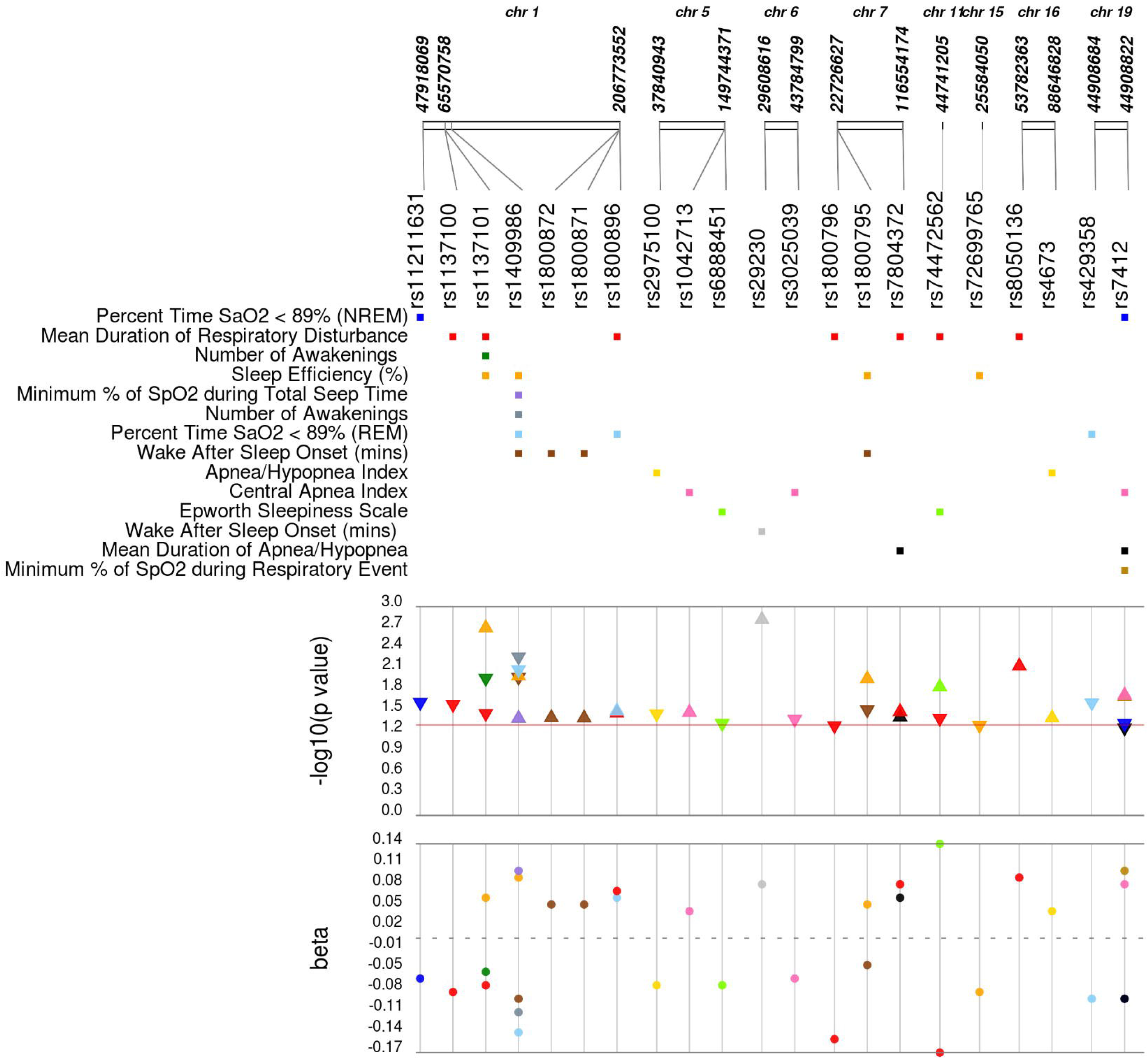
Associations between Literature-derived Candidate SNPs and Sleep Study Report Variables Available in Geisinger EHRs. Plotted are the estimates of the additive effects (beta) of the minor allele at each candidate SNP on Box-Cox transformed variables obtained from sleep study reports, along with the corresponding −log10 p-value. For p-values, up arrows denote increases in variable measurements and down arrows denote decreases.

As mentioned above, there were also 16 SNPs that were associated with quantitative sleep study report variables with no evidence for an association with EHR-OSA (Figure 2, Table S4). These included associations between a SNP in a non-coding transcript of *prostaglandin E receptor 3* (*PTGER3*) and two respiratory indices, as well as three measures of sleep efficiency (rs1409986, Table S4). Additional SNPs associated with more than one sleep study report trait included an intronic SNP in the *interleukin 6* (*IL6*) gene, rs1800795, which was associated with sleep efficiency (β=0.05, p=1.23×10^−2^) as well as wake after sleep onset (β=−0.04, p=2.67×10^−2^); an intronic variant in the *tetraspanin 18* (*TSPAN18*) gene, rs74472562, with the mean duration of a respiratory disturbance (β=−0.17, p=3.53×10^−2^) and reports of sleepiness (β=0.14, p=1.61×10^−2^); an intronic variant, rs7804372, in the *caveolin 1* (*CAV1*) gene with the mean duration of both a respiratory disturbance (β=0.08, p=3.65×10^−2^) as well as apnea/hypopnea (β=0.06, p=4.37×10^−2^); and a SNP, rs7412, tagging the ε2 allele in *APOE* with four different respiratory indices (Figure 2, Table S4).

### Phenome-wide association studies of OSA candidate variants

Seven of the SNPs previously reported in the literature to be associated with OSA were associated (q<5.0×10^−2^) with other clinical traits derived from the EHR in discovery analyses conducted in the Geisinger dataset (Figure 3, Tables 2 & S5-S6). Of these, four SNPs that were validated for an association with EHR-OSA in at least one analysis dataset were also associated with traits reflecting immune diseases, peripheral nerve disorders, diabetes and genitourinary symptoms (Figure 3, Tables 2, S3 & S6). Specifically, presence of the A allele at the intronic *LPAR1* variant, rs7030789, was associated with increased risk for EHR-OSA in the Geisinger dataset, as well as increased risk for evidence of nocturia (β=0.23, p=6.36×10^−6^, q=7.99×10^−3^; Figure 4a). The association with nocturia reflected in the PheCode for ‘Frequency of urination and polyuria’, but not EHR-OSA, generalized in the VUMC African American dataset (β=0.46, p=3.68×10^−2^, q=9.78×10^−1^). In addition, an association between the rs1800629 variant near *TNF-α* and celiac disease was observed in the entire Geisinger dataset (β=1.17, p=4.69×10^−23^, q=2.57×10^−19^); this SNP was primarily associated with EHR-OSA in individuals from Geisinger with BMI≤30 (β=−0.20, p=1.03×10^−2^). In this particular subgroup defined by lower BMI, the SNP near *TNF-α* was also associated with diabetes mellitus codes (β=0.09, p=4.09×10^−5^, q=1.47×10^−2^), and increased measures of blood glucose (β=2.08, p=6.98×10^−5^, q=1.68×10^−2^), hemoglobin A1c (β=0.06, p=1.67×10^−4^, q=3.01×10^−2^) and natriuretic peptide B (β=48.20 p=3.96×10^−5^, q=1.47×10^−2^). The association between the A allele at the rs1800629 SNP near *TNF-α* and evidence of celiac disease replicated in the entire VUMC European American dataset (β=1.03, p=1.30×10^−6^, q=6.11×10^−3^; Figure 4b). The same allele at this SNP was also associated with increased risk for EHR-OSA in this dataset (Table 2). In addition, an intronic *FTO* variant (rs8050136), while not associated with EHR-OSA in the entire Geisinger dataset, was associated with OSA severity reflected by a longer duration of respiratory disturbance and with diabetes mellitis (β=0.08, p=2.26×10^−6^, q=3.34×10^−3^). The association between the intronic *FTO* SNP and diabetes mellitis replicated in the VUMC European ancestral dataset (β=0.10, p=3.23×10^−3^, q=1.00). Notably, this SNP was associated with EHR-OSA specifically in males with European ancestry from VUMC and was also associated with diabetes mellitis in this same subgroup (β=0.09, p=3.02×10^−2^, q=9.99×10^−1^). Finally, the *NRG1* intronic rs10097555 variant which was only associated with EHR-OSA in the VUMC African American dataset (Figure 1, Table 2) was instead associated with evidence of peripheral nerve disorders in the Geisinger dataset (β=0.47, p=5.31×10^−6^, q=6.92×10^−3^); however, the association between this SNP and peripheral nerve disorders did not replicate in the VUMC European American dataset or generalize in the VUMC African American dataset (Table S5).

**Figure 3.**
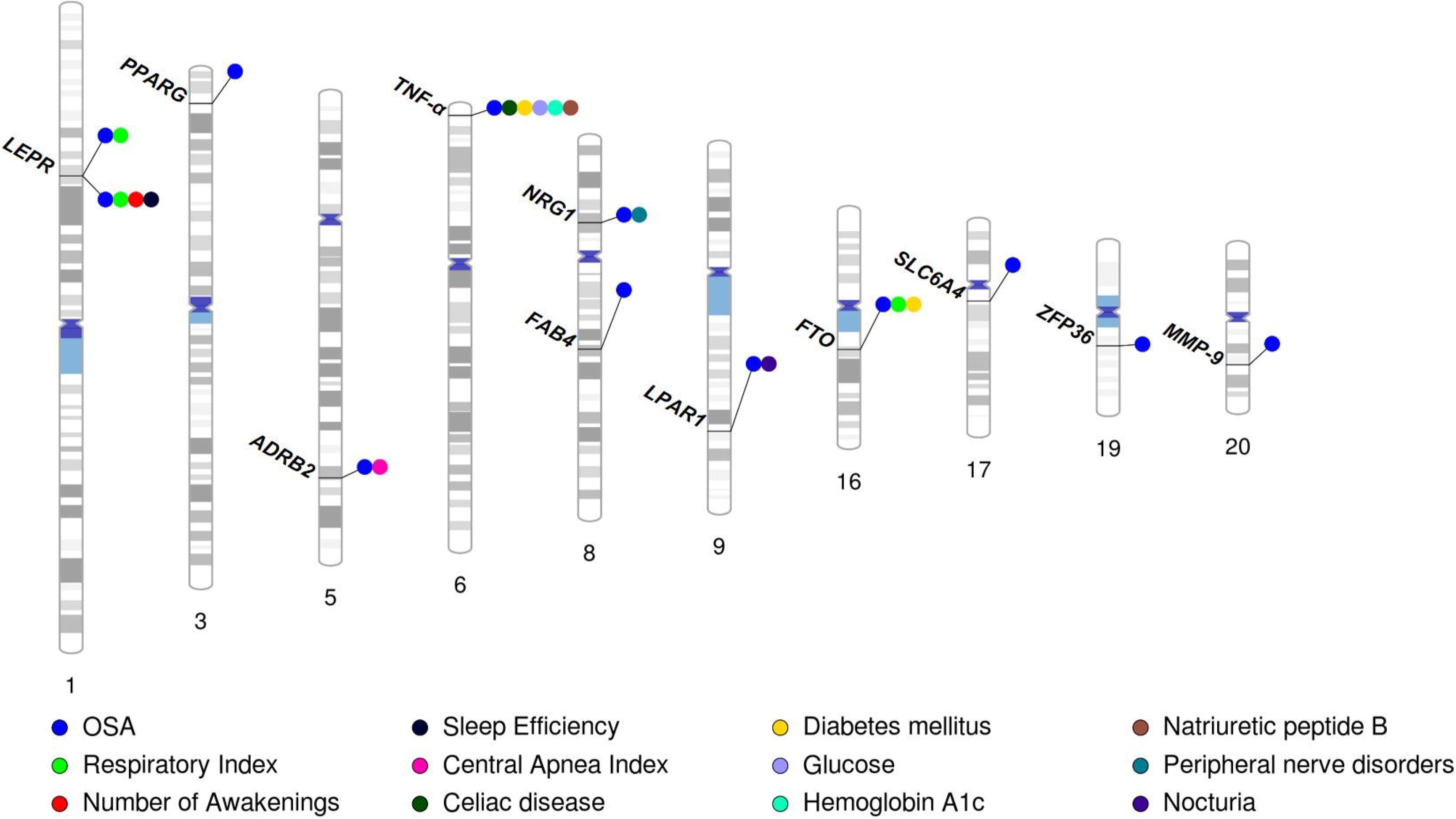
Sleep Report, Clinical and Laboratory Traits Associated with EHR-derived Obstructive Sleep Apnea-Associated SNPs. Phenogram plot of chromosomal locations for OSA candidate variants that were nominally associated (p<5.0×10^−2^) with EHR-derived OSA in at least one of the evaluated clinical datasets. Additional associations of these SNPs with traits obtained from EHRs reflecting variables from sleep reports, clinical codes and clinical laboratory values are shown. The abbreviation for the gene encoded in the genomic location closest to the SNP is noted.

**Figure 4.**
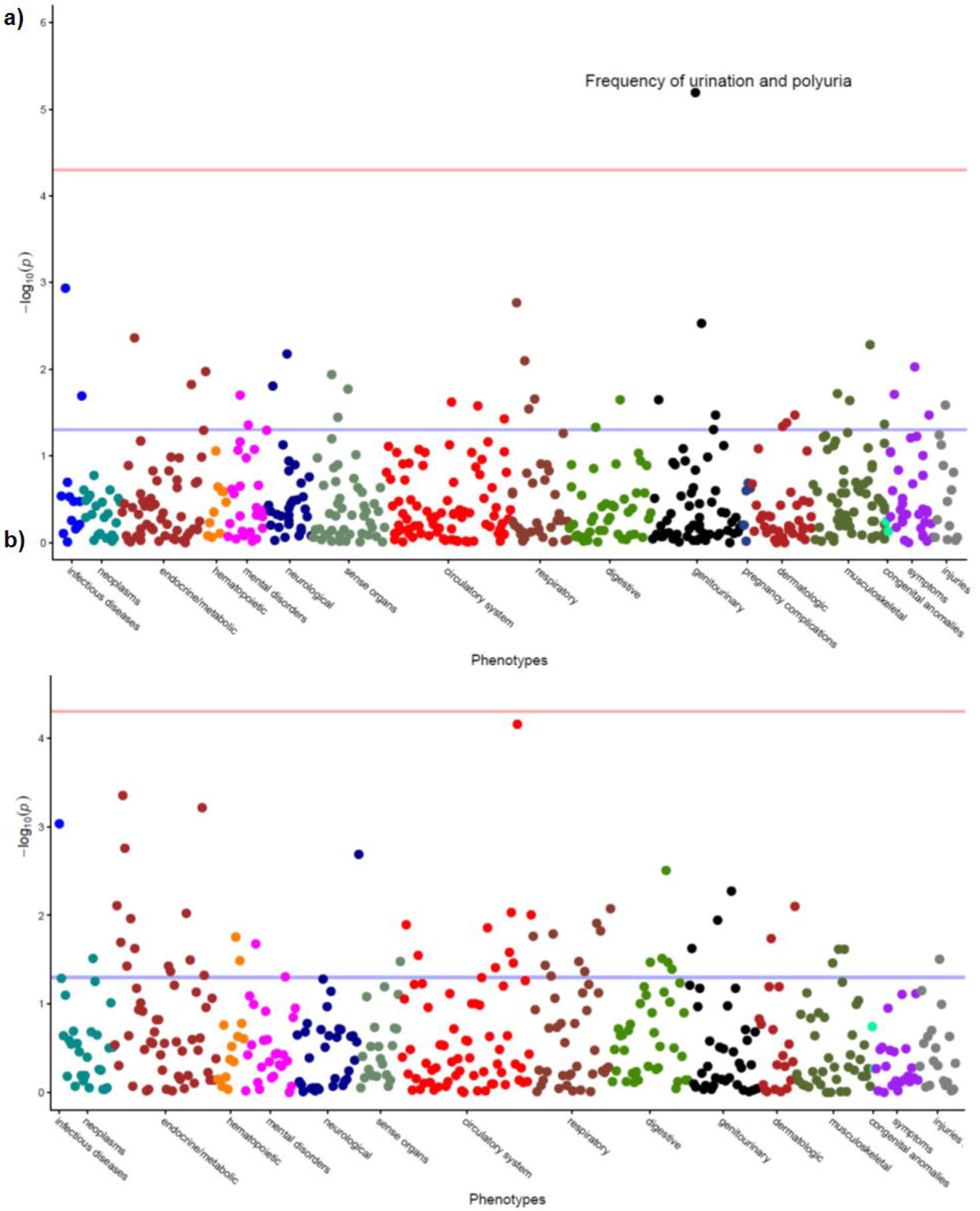
EHR-derived Clinical Traits Associated with EHR-OSA Hits. Manhattan plot of all phenotypes associated with the **a)** rs7030789 variant, which was also associated with EHR-derived OSA, in the Geisinger European American analysis dataset and the **b)** rs1800629 variant, which was also associated with EHR-derived OSA, in the VUMC European American analysis dataset. Red line denotes FDR-adjusted significance threshold (q<5.0×10^−2^), blue line denotes suggestive significance (p<5.0×10^−2^).

There were two OSA candidate SNPs that, while not associated with EHR-derived diagnosis of OSA in any of the datasets, were associated with measures of OSA severity defined using variables from sleep study reports (Table S4), as well as other clinical codes and laboratory values (Table S5). This included the SNP tagging the ε2 allele in *APOE* mentioned above and the ε4 allele tagging SNP (rs429358). Specifically, there were associations between the C allele at rs429358 and reduced time spent with oxygen saturation <89% during rapid-eye movement sleep (β=−0.09, p=2.12×10^−2^), as well as increased risk for hyperlipidemia (β=0.26, p=1.31×10^−23^, q=7.57×10^−20^), increased triglycerides (β=8.39, p=2.01×10^−15^, q=6.99×10^−12^), lower levels of HDL cholesterol (β=−1.51, p=1.73×10^−22^, q=8.80×10^−19^), and increased risk for coronary atherosclerosis (β=0.15, p=1.01×10^−5^, q=1.15×10^−2^). The associations between rs429358 in *APOE* with hyperlipidemia (β=0.16, p=1.06×10^−4^, q=2.28×10^−1^), and lower levels of HDL cholesterol (β=−6.12, p=5.94×10^−6^, q=3.15×10^−4^) replicated in the VUMC European American dataset; an association with hyperlipidemia (β=0.36, p=6.42×10^−4^, q=3.12×10^−1^) also generalized to VUMC African Americans. The rs7412 SNP was also associated with hyperlipidemia in both independent European American datasets (Table S5).

One OSA candidate SNP, rs2187668, an intronic variant in the *major histocompatibility complex, class II, DQ alpha 1* (*HLA-DQA1*) gene, had no evidence for an association with EHR-OSA or OSA severity but instead with codes for celiac disease in the Geisinger dataset (β=1.62, p=4.46×10^−42^, q=5.80×10^−38^), which replicated in the VUMC European American dataset (β=1.44, p=6.19×10^−12^, q=1.46×10^−7^). Additional associations between the *HLA-DQA1* SNP and autoimmune diseases of the thyroid (chronic thyroiditis: β=0.53, p=7.33×10^−6^, q=8.88×10^−3^; hypothyroidism: β=0.15, p=4.64×10^−7^, q=7.67×10^−4^) replicated in the VUMC European American dataset (chronic thyroiditis: β=0.51, p=1.17×10^−2^, q=1.00; hypothyroidism: β=0.19, p=1.31×10^−4^, q=2.37×10^−1^). While not replicated, rs2187668 was also associated with measures of calcium, glucose and hemoglobin A1c in the Geisinger dataset (Table S5).

## DISCUSSION

By harnessing the rich resources of clinical data made available from EHRs linked to large biorepositories at Geisinger and Vanderbilt University Medical Center, this study represents one of the largest and most comprehensive studies of the genetics underlying OSA and co-occurring conditions in OSA. In total, we observed associations between 29% (14/48) of the candidate OSA SNPs with EHR-derived definitions of OSA in at least one analysis dataset or covariate subgroup. Of these, 11 SNPs were observed to have effects in the same direction as those reported in the literature—four of which were also associated with OSA severity measures from sleep study reports. These variants may be particularly robust and more likely to translate into clinical populations treated in the United States. Furthermore, SNPs associated with EHR-OSA and/or OSA severity demonstrated evidence for pleiotropic effects on expression of other EHR-derived phenotypes, including celiac disease, diabetes, genitourinary symptoms, heart failure, and peripheral nerve disorders. SNPs which did not validate for an association with EHR-OSA were instead associated with celiac disease and autoimmune thyroid diseases. We expect that these results will help to tease apart the substantial heterogeneity underlying expression of OSA and provide the basis for more targeted, personalized treatment of OSA that can be tailored to particular symptom or comorbidity profiles.

### EHR-derived evidence supports influences of genetic variation in GABBR1 and LEPR on risk for OSA and OSA severity in United States clinical populations

Notably, the G allele at the rs29230 variant located in the *GABBR1* gene was associated with increased risk for EHR-OSA when results from analyses conducted separately in the Geisinger, VUMC European Americans and VUMC African Americans were meta-analyzed. This allele was also associated with longer wake after sleep onset in the subset of the Geisinger dataset with available sleep study reports. The association of the rs29230 variant and OSA was originally reported in a candidate gene study conducted in a relatively small population (n=174) of individuals from Turkey^61^. OSA was defined based on an AHI≥5 and controls were confirmed using medical histories. Notably, this study excluded individuals with evidence of an underlying metabolic disorder or chronic obstructive pulmonary disease. The authors observed an effect primarily in men indicating the C allele at rs29230 (reported in the literature on the alternative strand) was increased in OSA patients^61^. A subsequent candidate gene study conducted in 266 individuals from China confirmed an association between the C allele at rs29230 and risk for OSA defined as AHI≥5 in patients with no evidence of underlying confounding comorbid conditions, while additionally controlling for differences in BMI between cases and controls^62^. The *GABBR1* gene encodes a receptor for the main inhibitory neurotransmitter in humans, gamma-aminobutyric acid (GABA). GABA(B) receptors are expressed in hypoglossal motorneurons innervating the tongue, which are important for inhibiting tongue movement^62^. It is possible that variation in *GABBR1* influences risk for OSA by affecting activity of the encoded receptor resulting in dysfunction of the hypoglossal motoneurons^61; 62^. Notably, the functional consequences of the rs29230 missense variant on *GABBR1*, depending on the isoform, varies across numerous *in silico* prediction algorithms and is reported by some software to be possibly damaging and others benign (see Ensembl Variant Effect Predictor for the range of consequences, https://uswest.ensembl.org/index.html). Thus, additional functional characterization of this SNP would be useful to help elucidate its potential role in contributing to the expression of OSA.

There were also two missense variants in *LEPR* that had substantial evidence for a relationship with OSA in individuals with genetically informed European ancestry. Similar to the effects reported in previous studies^63^, the G alleles at the *LEPR* SNPs, rs1137100 and rs1137101, were associated with reduced risk for EHR-OSA in individuals from Geisinger. The same alleles at these SNPs were associated with shorter respiratory events measured via PSG. In addition, the rs1137101 SNP was associated with a reduced number of awakenings and increased sleep efficiency. These variants were originally evaluated in candidate gene studies as *LEPR* encodes a receptor for the adipocyte-specific leptin hormone and there is evidence of a relationship between leptin levels and OSA^64; 65^. Leptin has been shown to be an antioxidant molecule that could potentially counteract oxidative stress as a consequence of chronic intermittent hypoxia in OSA. Higher levels of this hormone were observed in OSA patients compared to controls^64^, and the AHI is an independent predictor of the evening/morning leptin ratio, suggesting that OSA might affect leptin diurnal rhythms^65^. Notably, in previously reported meta-analyses, the rs1137101 SNP was associated with reduced risk for OSA in European but not East Asian ancestries, while the rs1137100 SNP was evaluated solely in East Asian populations with no conclusive evidence for an association with OSA^63^. In particular, the original study reporting the European ancestral association between rs1137101 and OSA was conducted in a population from Poland with PSG-confirmed cases and controls matched for age, gender and BMI^66^. Given the strong relationship between OSA and obesity, it is interesting to note that these *LEPR* SNPs are reported in more than one study to be unassociated with risk for obesity^67–69^, suggesting variation in the *LEPR* gene may have effects on expression of OSA that are independent of obesity. It is unclear what are the functional consequences of these two missense mutations in *LEPR*; however, both are reported in ClinVar as benign (https://www.ncbi.nlm.nih.gov/clinvar/).

### Many OSA candidate SNPs associate with severity, not diagnosis

Of the 21 SNPs that were associated with respiratory indices and measures of sleep quality in the Geisinger dataset, there were 16 that were not associated with an EHR-derived definition of OSA in any analysis dataset or covariate subgroup. It is possible that these associations did not validate for a relationship with EHR-OSA because these SNPs have effects on OSA-related severity measures which are more robust than effects on expression of an EHR-derived OSA diagnosis, which relies on meeting International Classification of Sleep Disorders diagnostic criteria. Notably, one SNP validated specifically for an association with OSA severity similar to what was reported in the literature. The T allele at the intronic *TSPAN18* SNP, rs74472562, was associated with a shorter duration of the respiratory event in individuals with Latin American ancestry^70^, similar to what was observed in European Americans in our study. We also observed an association between this SNP and increased reports of sleepiness measured with the Epworth Sleepiness Scale. It is possible these results reflect a decreased arousal threshold and individuals with this variant are more likely to wake-up during an event which would explain why event durations are shorter and symptoms of sleepiness are increased.

Varying definitions of PSG-based OSA diagnoses within the reviewed literature could also reflect the associations with OSA severity, but not OSA diagnosis that we observed. For example, some studies defined OSA cases as having AHI≥5^71–73^, while others used AHI≥10^74^, AHI≥15^41; 44; 75^, or even presented varying definitions depending on the analysis^76^. Additionally, two reviewed studies were meta-analyses, with varying definitions of OSA depending on the selected study^77; 78^. Many previous studies reporting associations between the candidate SNPs we tested and OSA, compared cases to controls who were defined by these PSG-measured AHI cut-offs, but did not assess potential genetic effects on quantitative measures of OSA severity. It is also possible that even though our definition of EHR-OSA was validated with an excellent NPV, a proportion of EHR-defined non-cases have undiagnosed disease (see Limitations); this bias would be eliminated when relying on OSA severity measures from sleep studies.

Overall, the range of associations between OSA candidate SNPs and quantitative measures from sleep study reports highlight the variability of symptoms of patients who come to the clinic to be evaluated for an OSA diagnosis. For example, we observed an association between the reported OSA risk allele^77^ at the rs1800765 intronic SNP in *IL6* gene and increased sleep efficiency, as well as reduced wake after sleep onset. It is possible that these patients have OSA but, as they are not waking up following an event, are not reporting symptoms and are therefore not being captured in the clinical setting. In addition, the A allele at the rs1409986 *PTGER3* SNP was associated with the largest number of sleep study report variables, including an increased amount of time spent at low oxygen saturation levels during sleep, but a decreased number of awakenings and increased sleep efficiency. These individuals may have a particularly unfavorable combination of symptoms, as it is possible they have higher arousal thresholds and are therefore less likely to wake up during events. These patients are potentially not recognizing symptoms and as such may not come to the clinic for evaluation. Furthermore, a missense variant in *ADRB2* was associated with an increased Central Apnea Index in the Geisinger dataset, but not with EHR-OSA or the AHI. It was, however, associated with reduced risk for OSA diagnosis in the VUMC dataset indicating that the rs1042713 SNP is potentially associated with other forms of sleep disordered breathing. Finally, the *APOE* ԑ2 tagging SNP was associated with a number of measures of OSA severity and central apnea as well as numerous other comorbidities. These individuals may not be receiving a diagnosis of OSA due to a clinical focus on their other co-occurring conditions.

### Associations of OSA candidate SNPs may not generalize across populations, or may reflect underlying comorbidities

There were 19 SNPs (~40% of those tested) reported in the literature to be associated with OSA that did not show an association with either EHR-defined OSA or with measures of OSA severity in our study. Notably, a number of the associations between SNPs tested in our samples of European or African ancestry were identified in individuals representing populations with different ancestral backgrounds (e.g., Latin, East Asian). It is possible that the effects of these variants on OSA do not generalize to populations of European American or African American ancestry. Another potential reason that the associations between these candidate SNPs and EHR-OSA or OSA-related severity measures did not validate is that the original studies did not consider the possibility that the SNP was associated with a prevalent comorbidity in OSA, rather than OSA itself. There was one intronic SNP (rs2187668) in the *HLA* gene that instead of being associated with EHR-derived evidence for OSA, it was associated with autoimmune diseases of the thyroid and celiac disease, as well as median clinical laboratory values of calcium, glucose and hemoglobin A1c levels in plasma or serum. Studies suggest that hypothyroidism is a risk factor for OSA, with as many as 35% of individuals with these disorders having comorbid, or eventual OSA^79^. Potential mechanisms implicated in the development of OSA as a consequence of autoimmune thyroid diseases include deposition of mucoprotein in the upper airway, reduced neural output to airway musculature, abnormal ventilatory control, and a dual relationship with obesity^80^. In addition, a connection between OSA and celiac disease has been proposed^81^, described in more detail below. While the original study evaluating an association between the *HLA* variant and OSA excluded a number of important comorbidities that may confound results, there was no mention of excluding individuals with thyroid disorders or celiac disease^82^. As such, it is possible that a proportion of individuals included as cases had underlying autoimmune disorders that were contributing to expression of OSA. This could result in spurious associations in the original study and explain why our study did not observe a relationship between this *HLA* SNP and evidence of OSA in the EHR.

### Evidence for pleiotropic genetic effects of OSA candidate variants in LPAR1, FTO, TNF-α, and NRG1

A proportion of the candidate variants that were validated for an association with EHR-OSA or OSA severity overlap with those associated with risk for co-occurring conditions, suggesting shared etiologies. For example, variation in the *LPAR1* gene was associated with EHR-OSA and the frequency of urination, which could reflect an underlying renal disease. Notably, more severe OSA has been shown to contribute to a higher risk of kidney and pancreatic cancer^46^ and the encoded receptor is evidenced to play an important role in the development of kidney^83^ and pancreatic cancer^84^. Thus, pleiotropic effects of variation in the *LPAR1* gene may help to account for the relationship between more severe OSA and comorbid kidney or pancreatic cancer. In addition, a common variant, rs8050136, residing in the locus where *FTO* is encoded is considered to be one of the most robust obesity-associated loci^34^. A previous PheWAS was conducted to test for associations between rs8050136 and 1,645 ICD-9 codes obtained from EHRs available in the electronic Medical Records and Genomics (eMERGE) Network^85^. The rs8050136 variant was associated with codes for sleep apnea^85^. In the independent datasets we evaluated (all eMERGE Network samples were removed), the rs8050136 SNP was associated with EHR-OSA in males with European ancestry from VUMC, with longer durations of respiratory disturbance in the entire Geisinger analysis dataset, and codes for diabetes mellitus in European Americans from Geisinger and VUMC. These results provide evidence for pleiotropic effects influencing expression of obesity, diabetes and OSA in many individuals with European ancestry.

We also observed an association between the A allele at the rs1800629 SNP near *TNF-α* and increased risk for EHR-OSA in the VUMC European American dataset, consistent with effects reported in the literature (see Table S2). This allele also had a strong effect on the risk for celiac disease in both datasets. These results are also consistent with previously reported associations for celiac disease^86^. A connection between celiac disease and sleep disorders has been proposed^87^. Not surprisingly, symptoms of celiac disease (e.g. gastroesophageal reflux disease) may contribute to disturbed sleep; however, evidence suggests that sleep disorders in individuals with celiac disease are independent of gastrointestinal issues^87^. Furthermore, celiac disease and OSA share common features of lymphatic hyperplasia and local inflammation, and studies conducted in children report an increased prevalence of OSA in patients with celiac disease^81^. Notably, the direction of the effect for the association of the *TNF-α* SNP with EHR-OSA was different in individuals with lower BMIs at Geisinger, with no evidence for a relationship with celiac disease in this subgroup. It is possible that the effects of this variant on expression of both OSA and celiac disease relate to underlying issues with inflammation, mediated by obesity.

### Limitations and Future Directions

As mentioned above, a primary limitation of our EHR-based OSA phenotype is the likelihood of undiagnosed OSA among individuals defined as non-cases (e.g., those without OSA-related ICD-9/ICD-10 diagnosis codes). While prior studies among individuals referred for sleep studies suggest that a high percentage may have undiagnosed OSA^88–91^, these studies are typically biased given the increased risk among those already referred for diagnostic testing. In our previous validation study^55^, when calculating the likelihood of OSA based on the symptomless multi-variable prediction score^92^, almost 70% of non-cases had a predicted OSA probability <0.2, while fewer than 3% had a probability of 0.7 or above. Thus, there is strong evidence that a majority of individuals defined in our study as non-cases are true controls. Ultimately, bias introduced by including individuals with undiagnosed OSA in the comparison group of non-cases is expected to make it more difficult to identify significant associations and should not have as strong of an effect on the associations that were validated in this study. Further, we complement analyses testing for associations between candidate SNPs and OSA diagnoses with tests for associations between these SNPs and quantitative OSA severity measures obtained from sleep study reports. This should further account for potential bias related to inaccurate definitions of control status which could make validation more difficult. Regardless, future OSA research using EHR datasets would benefit from studies focused on expanding algorithms to more accurately identify undiagnosed cases of OSA in EHR-derived control samples.

Other limitations include the possibility that differing criteria for sleep apnea (hypopneas) at each site influenced results. Specifically, the AHI is the primary measure used to define the diagnostic criteria for OSA. The criteria for what is considered a hypopnea—which is instrumental to the calculation of the overall AHI—has changed within the last decade^93^. As such, use of different scoring criteria for hypopneas results in differential diagnoses of OSA. The possibility that the two clinical sites included in our studies may have adhered to different criteria for hypopnea definitions at various times throughout the duration of the EHR data we analyzed may account for why we observed unique associations between OSA candidate SNPs and EHR-derived OSA when comparing Geisinger to VUMC, particularly with regard to European Americans. However, given that we excluded individuals with only one diagnostic code on different dates at VUMC, and with two codes at Geisinger, we do not expect that this substantially influenced our results. Future work aimed at understanding the proportion of patients who fail to be diagnosed following a sleep study due to differences in hypopnea scoring criteria would be beneficial to alleviating this potential limitation to using EHR-derived datasets.

Overall, by capitalizing on the breadth of available clinical information in the EHR, we confirmed associations between many previously reported OSA candidate SNPs and OSA diagnosis, as well as OSA severity. Furthermore, we helped to establish genetic relationships between OSA and other diseases providing evidence for pleiotropic genetic effects influencing expression of multiple disorders in the same individual. Our results reflect the remarkable resources that EHR-derived datasets offer which are useful to characterizing genetic and phenotypic heterogeneity for complex human disorders, like OSA.

## Supplemental Data

Supplemental data includes six tables and two supplemental figures.

## Acknowledgements

This work was supported by the following grants from the National Institutes of Health: NHLBI R01HL134015 (AIP, JDR, SAP, KB, BAM), NLM K01LM012870 (OJV), NHLBI P01HL0943 (AIP), AASM Foundation 194-SR-18 (DRM). We acknowledge the resources provided by the Biorepository of Vanderbilt University (BioVU) team, specifically Jonathan T. Fleming, Celestial R. Jones-Paris and Lana M Olsen. Vanderbilt University’s BioVU projects are supported by numerous sources: institutional funding, private agencies, and federal grants. These include the NIH funded Shared Instrumentation Grant S10RR025141; CTSA grants UL1TR002243, UL1TR000445, and UL1RR024975. Genomic data are also supported by investigator-led projects that include U01HG004798, R01NS032830, RC2GM092618, P50GM115305, U01HG006378, U19HL065962, R01HD074711; and the following additional funding sources (https://victr.vanderbilt.edu/pub/biovu/?sid=229). This project was also funded by a grant from the Pennsylvania Department of Health (#SAP4100070267). The Department of Health specifically disclaims responsibility for any analyses, interpretations, or conclusions.

