## Supplemental Figures for "Characterization of Genetic and Phenotypic Heterogeneity of Obstructive Sleep Apnea Using Electronic Health Records"

**Figure S1. Inclusion Criteria for Studies Reviewed for Selection of Candidate Variants.**

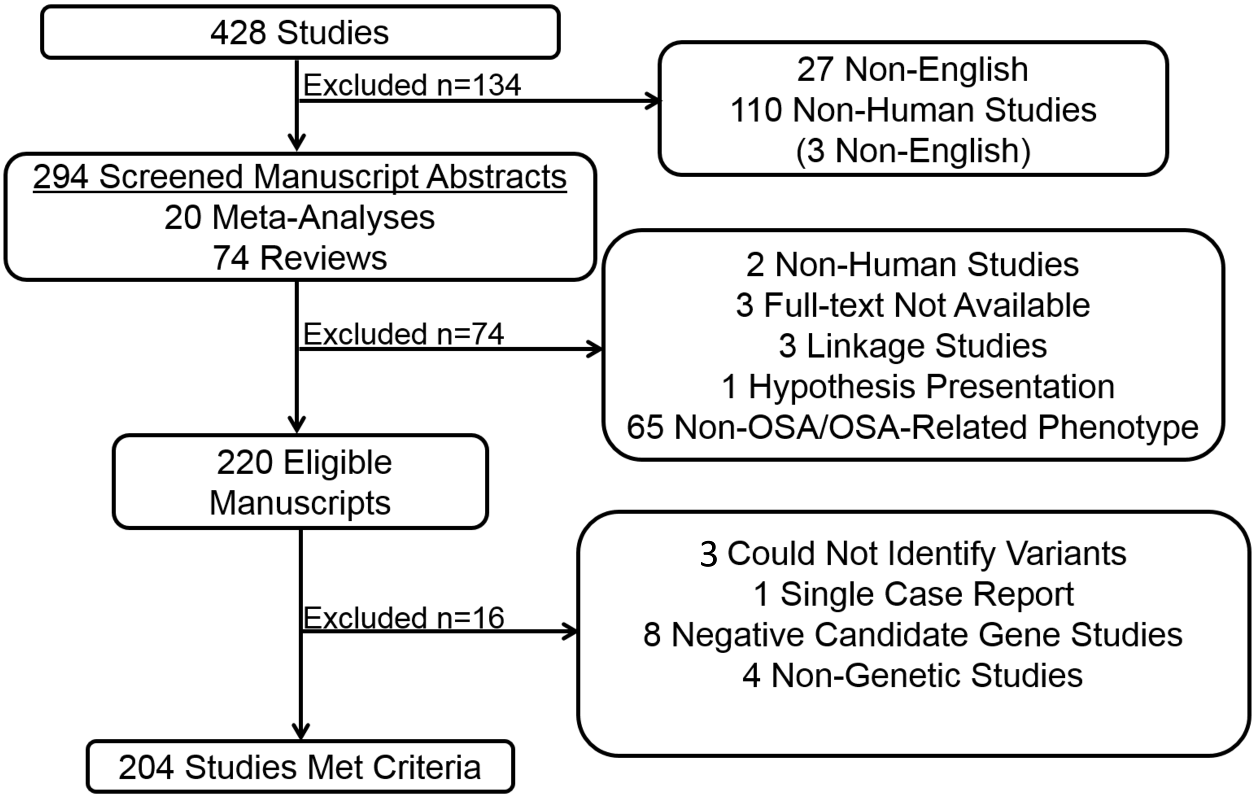

**Figure S2. Associations between Literature-derived Candidate SNPs and EHR-derived Obstructive Sleep Apnea in Subgroups Defined by Age, Sex, and BMI.** Plotted are the –log10 p-values of tests for associations between SNPs where interaction tests indicated an effect of Age, Sex, and/or BMI, and OSA defined in the EHR conducted in the entire dataset and separately in covariate-defined subgroups. Geisinger European American (EA) dataset results are plotted in blue, VUMC European Americans (EA) in red and VUMC African Americans (AA) in green. Up arrows denote increased risk for EHR-derived OSA given the minor allele at this SNP and down arrows denote reduced risk.

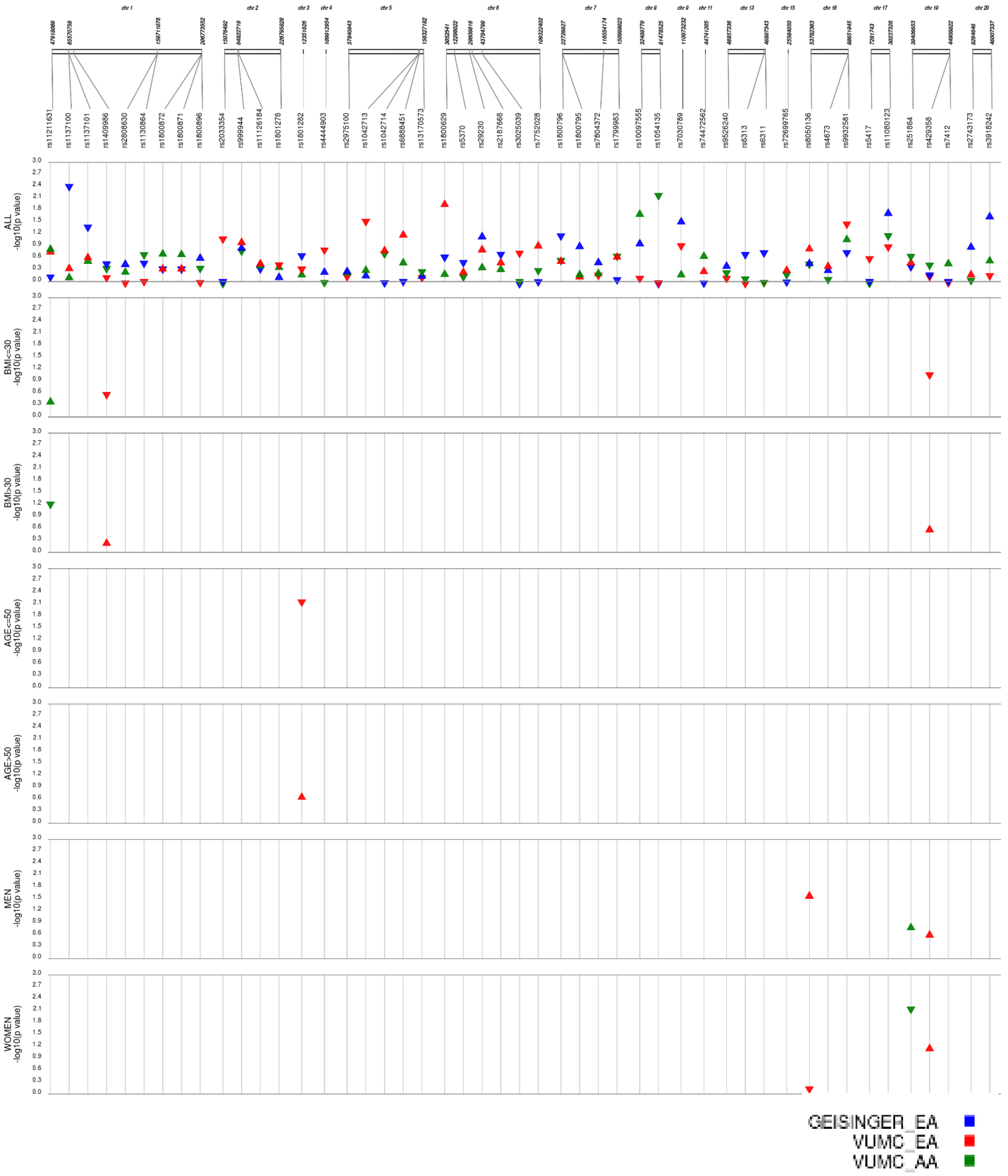
